# ECM-integrin signalling instructs cellular position-sensing to pattern the early mouse embryo

**DOI:** 10.1101/2021.07.05.450860

**Authors:** Esther Jeong Yoon Kim, Lydia Sorokin, Takashi Hiiragi

## Abstract

Development entails patterned emergence of diverse cell types within the embryo. In mammals, cells positioned inside the embryo gives rise to the inner cell mass (ICM) that eventually forms the embryo proper. Yet the molecular basis of how these cells recognise their ‘inside’ position to instruct their fate is unknown. Here we show that cells perceive their position through extracellular matrix (ECM) and integrin-mediated adhesion. Provision of ECM to isolated embryonic cells induces ICM specification and alters subsequent spatial arrangement between epiblast (EPI) and primitive endoderm (PrE) cells that emerge within the ICM. Notably, this effect is dependent on integrin β1 activity and involves apical to basal conversion of cell polarity. We demonstrate that ECM-integrin activity is sufficient for ‘inside’ positional signalling and it is required for proper sorting of EPI/PrE cells. Our findings thus highlight the significance of ECM-integrin adhesion in enabling position-sensing by cells to achieve tissue patterning.

## Introduction

Development begets an immense diversity of animal forms as fertilisation is followed by organisation of cells into higher order structures. The emergence of complex patterns generally requires that cells continuously exchange signals with their surroundings to direct their fate and spatial orientation. While transcriptional networks inform individual cell type, the *position* at which a cell lies is critical for tissue patterning. Therefore, relay of spatial information is a ubiquitous requirement in developing systems, and a cell has to sense its position relative to its neighbours to support robust patterning. A cell may glean positional information from a variety of sources, such as mechanochemical gradients, wave-like propagation of signalling activity, as well as direct adhesive interactions with the immediate environment (Jouve et al., 2002; Steinberg and Poole, 1981; Wolpert, 1969).

In particular, adhesive interactions with the extracellular matrix (ECM) are dynamically engaged in development and homeostatic turnover of tissues (Gattazzo et al., 2014; Walma and K. M. Yamada, 2020). The ECM consists of a network of various components such as laminin, collagen IV, and fibronectin, which serve to regulate cell behaviours ranging migration, polarisation, survival, and differentiation. Its significance is evident during development, where loss of laminin chains, collagen IV, or their respective receptors leads to early embryonic lethality in mice (Miner et al., 1998; 2004; Smyth et al., 1999; Williamson et al., 1997). Furthermore, laminin regulates gene expression and spatial organisation of cells in several epithelial tissues (Klein et al., 1988; Streuli et al., 1995). In the gut, its loss leads to epithelial hyperplasia and an impaired stem cell pool, while provision of ECM through Matrigel supports long-term culture of intestinal crypt organoids (Fields et al., 2019; Sato et al., 2009). Similarly, laminin is required for proper positioning and maintenance of follicle stem cells in their niche within the *Drosophila* ovary (O’Reilly et al., 2008). As such, laminin as well as other ECM components have a conserved role in modulating the spatial organisation and behaviour of cells across diverse contexts.

The preimplantation mouse embryo is remarkable in its regulative capacity to preserve embryonic patterning against drastic reduction in cell number (Solter and Knowles, 1975; Tarkowski, 1959; Tarkowski and Wróblewska, 1967). This implies dynamic readout of positional information by blastomeres to adjust their fate and spatial arrangement in response to perturbations. By the end of the preimplantation stage at embryonic day (E) 4.5, the embryo consists of an outermost trophectoderm (TE) monolayer enclosing a fluid-filled cavity and an inner cell mass (ICM). Within the ICM, the epithelial primitive endoderm (PrE) lines the cavity while epiblast (EPI) cells reside between the PrE and the overlying TE.

Cell position instructs the first lineage segregation in mouse development, as inner and outer cells become ICM and TE, respectively (Rossant and Tam, 2009; Tarkowski and Wróblewska, 1967). Prior to TE specification, the outer surface of the embryo is marked by a polarised cortical domain enriched in phosphorylated ezrin, radixin, moesin (pERM), Par6 and atypical protein kinase C (aPKC) (Ducibella et al., 1977; Louvet et al., 1996; Vinot et al., 2005; Ziomek and Johnson, 1980). This apical domain is both necessary and sufficient for TE fate, and effectively serves as the ‘outside’ positional signal to prompt embryonic patterning (Alarcon, 2010; Korotkevich et al., 2017). Conversely however, insights into ICM development and its underlying ‘inside’ cues thus far remain sparse. Despite advances made with embryonic and induced pluripotent stem cells, there is meagre understanding of how cells with the potential to form the entire adult animal actually arise *in vivo* (Evans and Kaufman, 1981; Takahashi and Yamanaka, 2006). Although the ICM gives rise to the embryo proper, its study is hampered by limited accessibility and the fact that early blastomeres tend to default to a TE-like state in response to various experimental perturbations (Korotkevich et al., 2017; Lorthongpanich et al., 2012; Stephenson et al., 2010; Tarkowski and Wróblewska, 1967).

## Results

### Inside positioning drives ICM specification during preimplantation mouse development

The earliest marker of ICM specification is the expression of *Sox2* (Guo et al., 2010; Wicklow et al., 2014). Therefore, we imaged *Sox2*-GFP x membrane tomato (mT) embryos with an inverted light-sheet microscope at 15-minute intervals to monitor the emergence of inside-outside cells and subsequent ICM specification (Figure 1A) (Arnold et al., 2011; Muzumdar et al., 2007; Strnad et al., 2016). This setup allowed long term live-imaging of 8-cell stage embryos until the early blastocyst stage with minimal phototoxicity. Given the natural embryo-to-embryo variability in developmental timing, embryonic age was standardised at the 8-to-9 cell division. Dynamic cell movements and asynchronous timing of division within each embryo led to internalisation of cells at different time points, consistent with earlier studies (Movie S1 and S2) (Dietrich and Hiiragi, 2007; Fleming, 1987). *Sox2*-GFP signals emerged only in completely internalised cells, and progressively increased in intensity as the embryo matured (Figure 1B and S1A).

**Figure 1.**
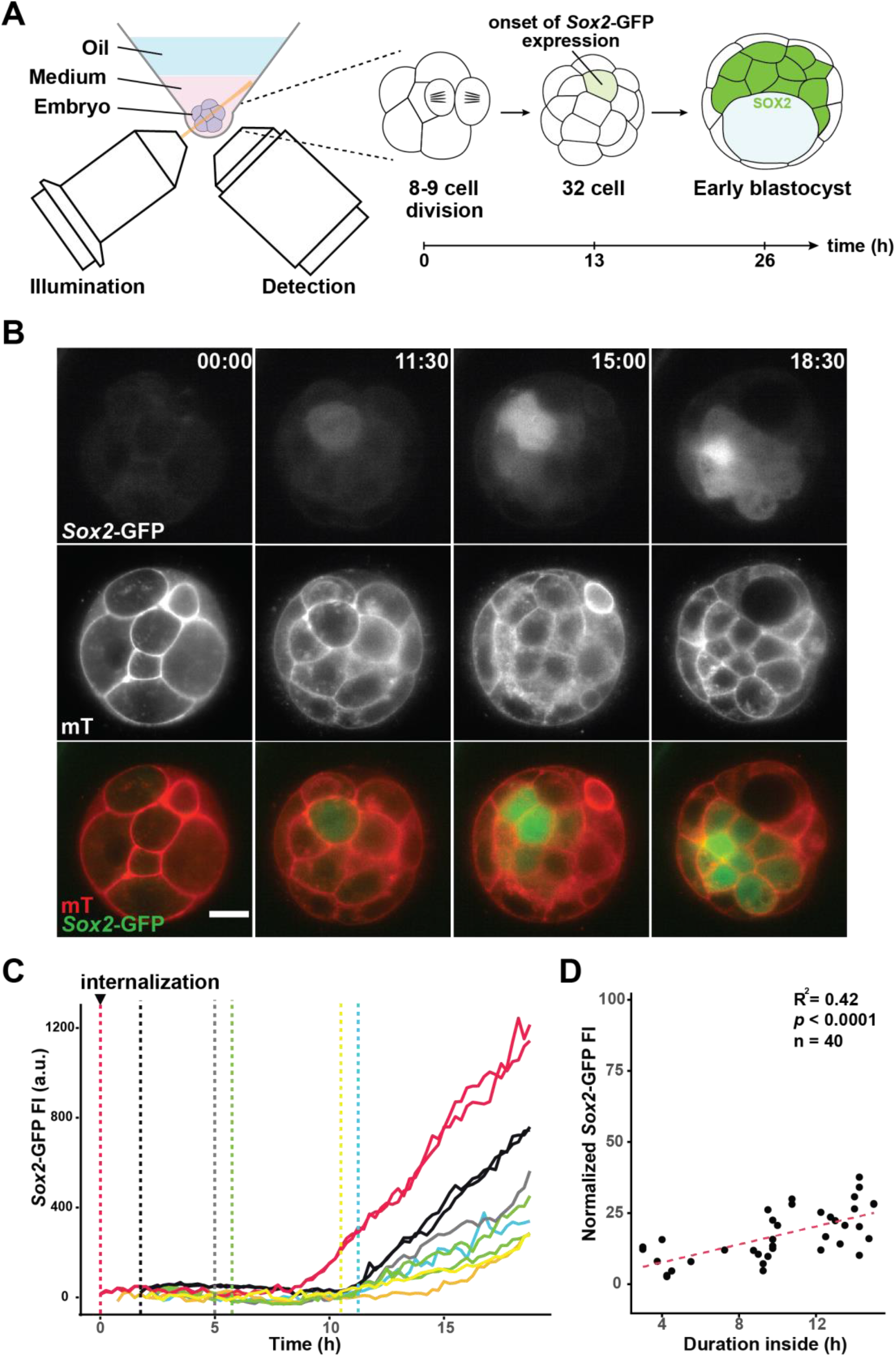
ICM fate specification is induced by inner positioning of the blastomere. (A) Schematic representation of live-imaging setup for preimplantation mouse development using selective plane illumination microscopy (SPIM). Fluorescent embryos expressing *Sox2*-GFP and membrane tomato (mT) were imaged from the 8-cell stage until the early blastocyst stage at 15-minute intervals. For subsequent image analysis, time (t) = 0 was set at the 8-to-9 cell division. (B) Time-lapse images of ICM specification during preimplantation development of a representative *Sox2*-GFP x mT embryo. Time, post 8-to-9 cell division (h:min). Scale bar = 20 μm. (C) Changes in *Sox2*-GFP intensity in inner cells during ICM specification. Cells become internalised at different timepoints (vertical dashed lines). Each line represents a single cell that becomes ICM-specified, and sister cells are marked in the same colour. Plot shows retrospectively tracked fluorescence intensities of 10 ICM cells from a representative embryo. (D) Scatterplot of normalised *Sox2*-GFP intensity and duration a cell resides in the embryonic interior completely surrounded by neighbouring cells. For each embryo, *Sox2*-GFP intensity was normalised to mean levels measured in the blastocyst. Pearson’s correlation was calculated at t = 15 h when heterogeneity in *Sox2* expression becomes apparent across different embryos. N = 4 embryos, n = 40 cells. See also Figure S1.

To follow changes in *Sox2* expression during ICM specification, ICM cells in the early blastocyst were retrospectively tracked to measure GFP fluorescence intensity at each time point (Figure 1C and S1B). Tracking extended to parent cells, noting the point at which each cell or its parent became internalised within the embryo. This revealed that *Sox2* expression levels during ICM specification is positively correlated with early internalisation (Figure 1D). These demonstrate that inside positioning of the blastomere is crucial for ICM specification

### Integrin and laminin chains are localised at the cell-cell interface

To identify the ICM-inducing cues of the embryonic intrior, we examined proteins enriched at the cell-cell interface. E-cadherin is clearly localised to cell-cell contact sites from the morula to blastocyst stages, away from TE-associated apical domains enriched in pERM (Figure 2A and 2B). While E-cadherin is the major adhesive molecule that holds the cells together irrespective of their fate (Filimonow et al., 2019; Larue et al., 1994; Shirayoshi et al., 1983; Stephenson et al., 2010), several studies have shown that ECM components are also present during this period (Dziadek and Timpl, 1985; Leivo et al., 1980; Morin and Sullivan, 1994; Sutherland et al., 1993).

**Figure 2.**
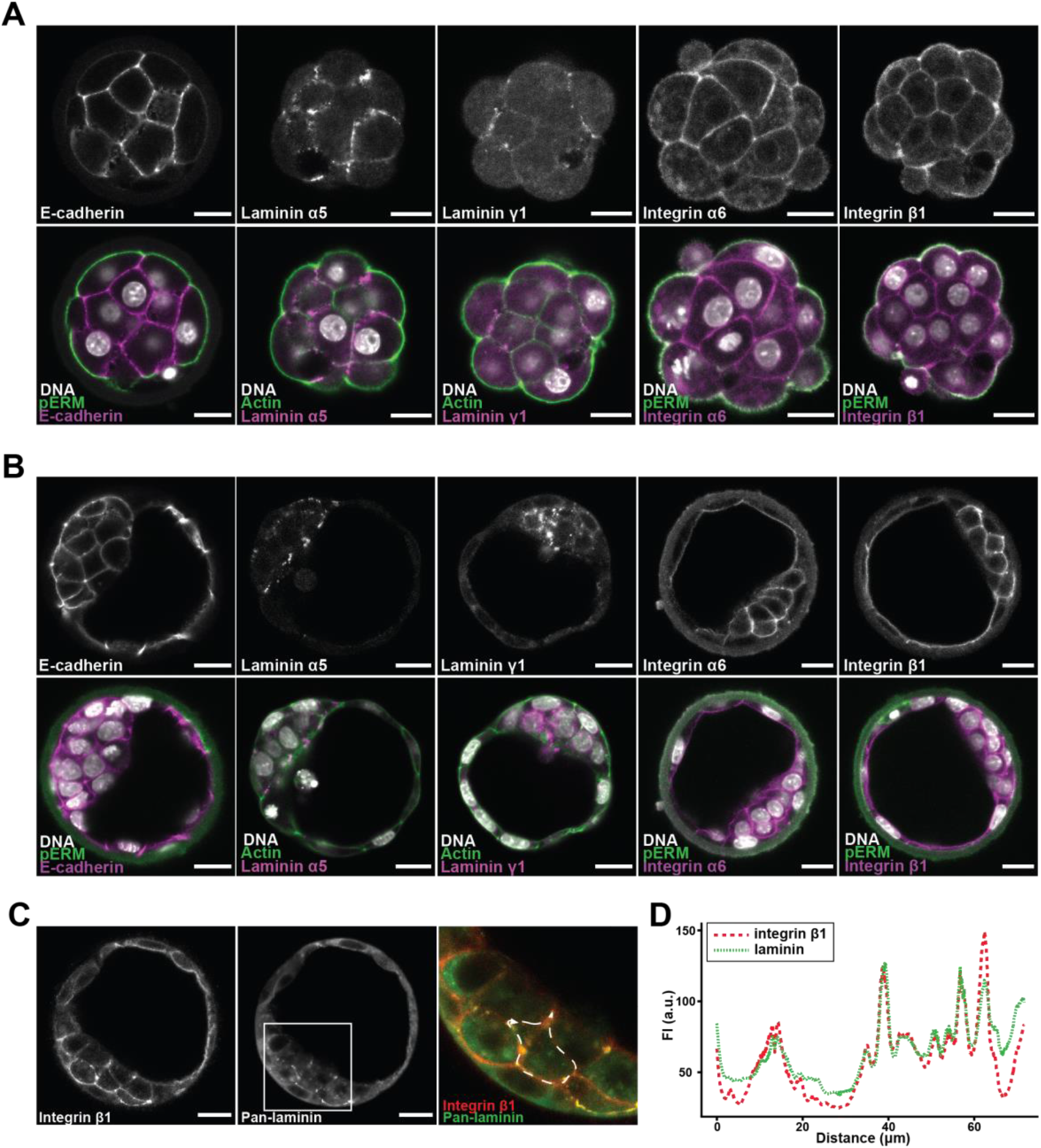
Integrin and laminin are expressed in the morula and colocalise in the blastocyst. (A and B) Localisation of E-cadherin, laminin chains α5 and γ1, and integrin α6 and β1 subunits in morulae (A) and blastocysts (B). Phosphorylated ezrin, radixin, moesin (pERM) marks the cell-free apical surface of outer cells. (C) Co-immunostaining for integrin β1 and laminin (non-chain-specific) in the blastocyst marks their shared localisation at the cell-cell interface. Scale bars = 20 μm. (D) Representative intensity profile of integrin β1 and laminin around an inner cell (marked by dashed white line in (C), arrowhead indicates starting point of measurement).

We found that several laminin chains are enriched at the cell-cell interface in the morula and the ICM region of the blastocyst (Figure 2A and 2B). Immunostaining indicated expression of laminin 511 in addition to the already reported laminin 111, which are heterotrimers of constituent α5, β1, γ1 and α1, β1, γ1 chains, respectively (Cooper and MacQueen, 1983; Leivo et al., 1980; Miner et al., 2004; Smyth et al., 1999). Accordingly, subunits of the major laminin receptor, integrin α6β1, which bind both laminin 111 and 511, were similarly expressed in the preimplantation embryo (Figure 2A and 2B) (Sutherland et al., 1993; Takizawa et al., 2017; M. Yamada and Sekiguchi, 2015). Close spatial correlation of laminin and integrin β1 fluorescence around inner cells identified ECM-integrin interactions as candidate ‘inside’ positional signals to blastomeres that drive ICM specification (Figure 2C and 2D).

### Exogenous ECM drives ICM specification and surface integrin α6β1 enrichment

To test whether the ECM provides ‘inside’ positional signals to drive ICM specification, we sought to mimic the inner environment of the embryo by providing the ECM through Matrigel, which is rich in laminin 111 (Orkin et al., 1977; Timpl et al., 1979). Embryos were recovered at the morula stage prior to marked upregulation of *Sox2*, and TE-specified outer cells were removed by immunosurgery (Figure 3A). Immunosurgery not only isolates naïve inner cells, but also alters positional identity by exposing them to the external environment (Solter and Knowles, 1975). Subsequent culture of these cells in standard embryo media (KSOM) fully restored inside-outside patterning. CDX2-positive TE cells surrounded SOX2-positive ICM cells and often a small fluid-filled cavity, reminiscent of blastocysts (Figure 3B, top panel). In this way, these isolated cells displayed robust regulative capacity.

**Figure 3.**
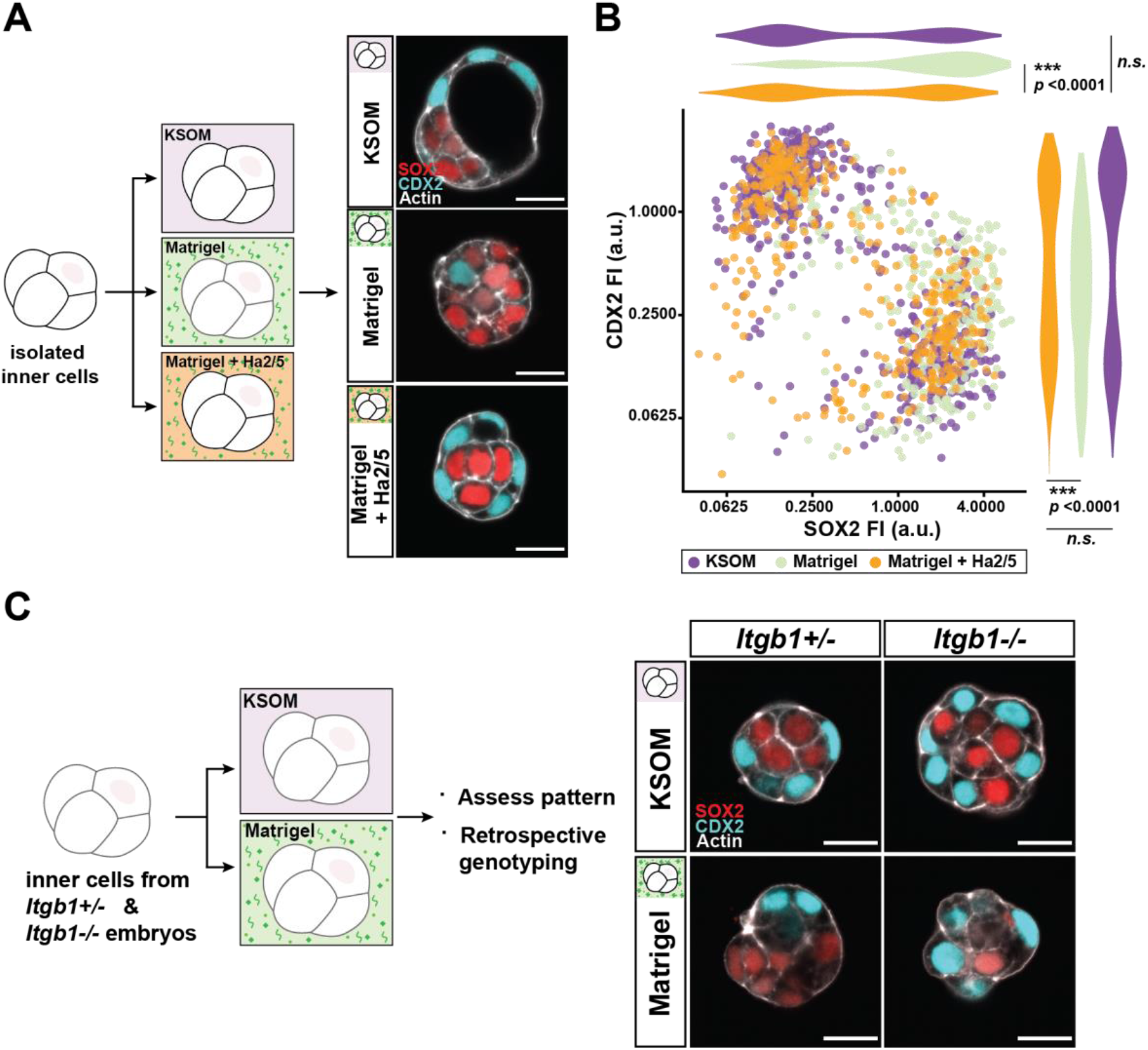
Exogenous ECM drives ICM specification and surface integrin α6β1 enrichment. (A) Schematic representation of experimental conditions and immunosurgery. Morula stage embryos are recovered prior to ICM specification, and lysis of outer cells leaves behind isolated inner cells. Immediately following immunosurgery, inner cells are cultured in either standard KSOM mouse embryo media or Matrigel before assessment of patterning. (B) Representative images of TE-ICM fate specification following immunosurgery and culture in either control KSOM or Matrigel. CDX2 marks TE fate while SOX2 marks ICM fate. In a few cases, Matrigel culture induces SOX2 upregulation across the entire cell cluster (bottom panel). Scale bars = 20 μm. (C) Total cell count after immunosurgery and culture in either KSOM (purple) or Matrigel (green). Each data point represents inner cells cultured from a single embryo. Student’s *t*-test, two-sided. Error bars show mean ± s.d. N = 43 embryos. (D) Scatterplot and adjacent violin plots show normalised fluorescence intensities of CDX2 and SOX2 measured for each cell cultured in either KSOM (purple) or Matrigel (green). Mann-Whitney U test. N = 43 embryos (n = 912 cells). (E and F) Representative images of pERM (apical marker), integrin β1, and integrin α6 localisation in cultured inner cells. (G and H) Quantification of surface enrichment of integrin β1 and pERM fluorescence. Student’s *t*-test, two-sided. Error bars show mean ± s.d. N = 49 (F) and N = 38 (G) embryos. (I) Circularity as a descriptor of cell shape measured for individual TE- and ICM-specified cells across the two culture conditions. Student’s *t*-test, two-sided. Error bars show mean ± s.d. N = 46 embryos (n = 288 cells). See also Figure S2.

In stark contrast, however, the TE layer was not restored in the presence of Matrigel. Instead, isolated cells formed a compact mass where the majority of nuclei were SOX2-positive, irrespective of cell position (Figure 3B). CDX2-positive cells were fewer and clustered at the periphery, while fluid-filled cavities were noticeably absent. Moreover, samples entirely composed of SOX2-positive cells were also observed across independent experiments, albeit at low frequency (9 out of 97, 9.3 %) (Figure 3B, bottom panel). Total cell numbers were comparable between the two conditions (Figure 3C), indicating that Matrigel does not have adverse effects on cell survival or proliferation.

Besides expression of *Cdx2* and *Sox2*, TE and ICM cells are distinguishable by differential Hippo signalling (Nishioka et al., 2009; Wicklow et al., 2014). In inner cells, Hippo signalling results in phosphorylation and cytoplasmic retention of YAP. In outer cells, Hippo signalling is inactive, and YAP translocates to the nucleus to drive downstream transcription of *Cdx2*. Consistent with increased *Sox2* expression, nuclear YAP localisation was diminished in Matrigel culture (Figure S2A). Furthermore, quantitative analysis of individual nucleus for levels of each fate marker confirmed significant increase in *Sox2* expression in Matrigel (Figure 3D). These findings demonstrate that exogenously supplied ECM provides ‘inside’ positional cues sufficient to drive ICM specification following immunosurgery.

Earlier studies noted that TE specification is preceded by ready polarisation of the outer surface after perturbations such as immunosurgery (Stephenson et al., 2010; Wigger et al., 2017). In agreement to this, pERM was enriched on the outer surface of isolated cells upon culture in KSOM, while integrin β1 was limited to cell-cell interfaces (Figure 3E, top panels). Distinct and mutually exclusive localisation of pERM and integrin β1 is consistent with the apicobasal polarity that accompanies inside-outside patterning in the whole embryo. In contrast, however, Matrigel led to significant enrichment of integrin β1 on the outer surface whereas peripheral pERM was significantly diminished (Figure 3E, bottom panels, 3G and 3H). Discontinuous patches of pERM were sometimes present on the surface, which generally coincided with CDX2-positive or SOX2-negative nuclei (Figure S2B). Integrin α6 localisation was comparable to integrin β1 (Figure 3F), while E-cadherin was limited to cell-cell interfaces regardless of culture conditions (Figure S2C). These suggests that Matrigel, particularly its constituent laminin, brings the integrin α6β1 receptor to the surface in lieu of apical polarity proteins, befitting ‘inside’ cells.

While ICM cells are roughly isotropic in shape, TE cells are generally oblong as these are stretched around the ICM or the fluid cavity (Chan et al., 2019; Niwayama et al., 2019). However, Matrigel abrogated this difference in circularity between TE and ICM cells. The presence of round TE cells in exogenous ECM suggests that fate specification here is not dependent on cell shape (Figure 3I).

### Integrin β1 activity is required for ECM-induced ICM specification

To test whether the activity of surface enriched integrin α6β1 is actually required for ICM induction by Matrigel, integrin β1 was inhibited with a function-blocking antibody, Ha2/5 (Mendrick and Kelly, 1993). Administration of Ha2/5 almost completely attenuated the effects of Matrigel. Despite the presence of exogenous ECM, outer cells polarised and became TE specified, while ICM specification was confined to inner cells (Figure 4A, 4B and S3A). In this way, cells cultured in Matrigel with Ha2/5 were indistinguishable from control samples. Similar observations were made upon inhibition of integrin α6 and assessment of YAP localisation (Figure S3B and C) (Sonnenberg et al., 1987).

**Figure 4.**
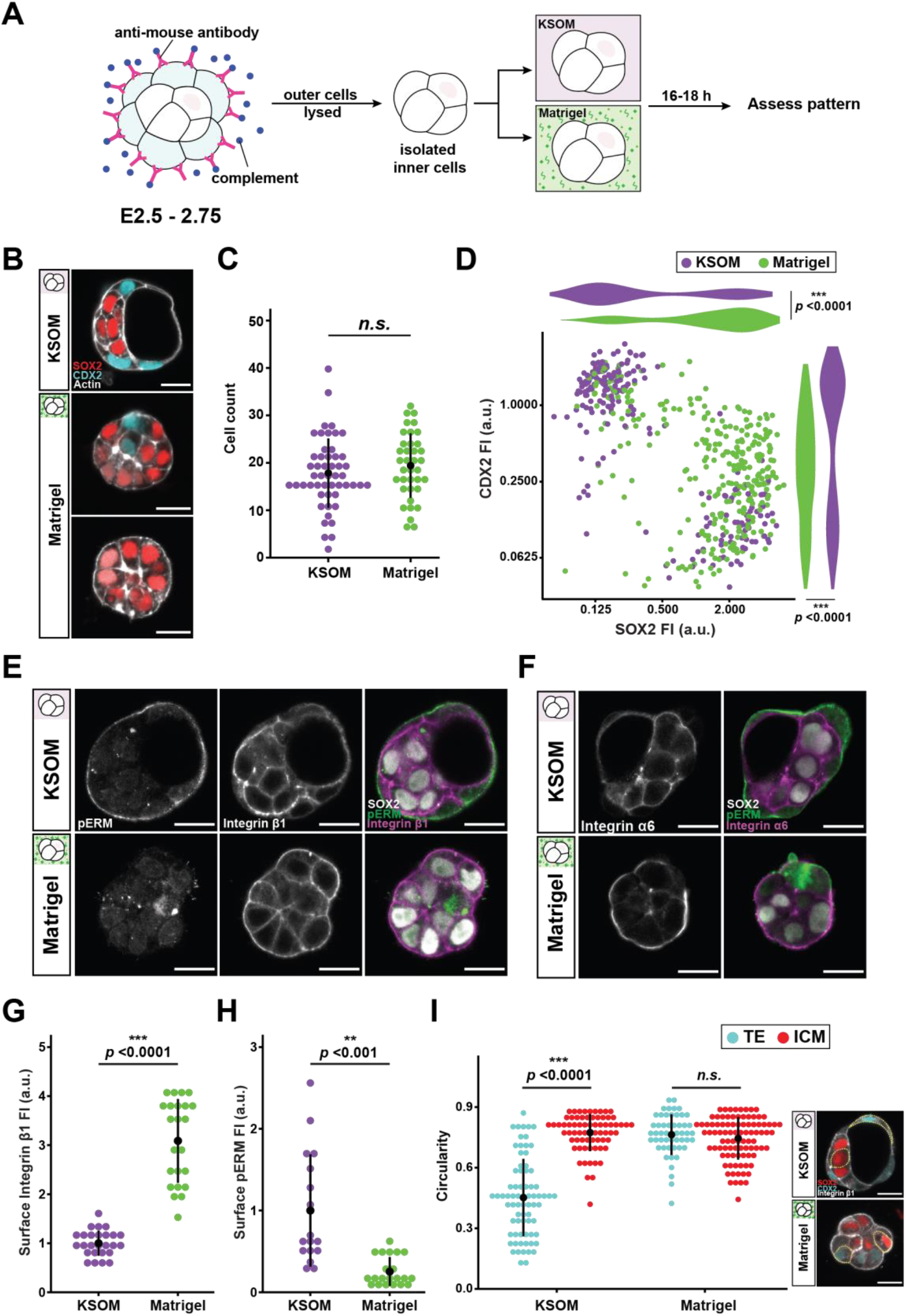
ICM induction by Matrigel is dependent on integrin β1 activity. (A) Schematic representation of experimental conditions and representative images of TE-ICM patterning upon administration of integrin β1 function-blocking antibody, Ha2/5 (10μg/ml) with Matrigel. (B) Scatterplot and adjacent violin plots show normalised fluorescence intensities of CDX2 and SOX2 measured for each cell cultured in either KSOM (purple), Matrigel only (light green) or Matrigel with Ha2/5 (orange). Mann-Whitney U test. Plot represents data combined from N = 93 embryos (n= 1332 cells). Data for KSOM and Matrigel are duplicated from Figure 3D for ease of comparision. (C) Schematic representation of experimental conditions and representative images of inner cells isolated from *Itgb1* transgenic embryos cultured in either KSOM or Matrigel. Each sample is genotyped retrospectively to identify *Itgb1*^-/-^ samples. N = 25 embryos (14 *Itgb1*^+/-^ and 11 *Itgb1*^-/-^). Scale bars = 20 μm. See also Figure S3.

Furthermore, upon genetic ablation of *Itgb1*, integrin β1-deficient cells were refractory to the effects of Matrigel (Raghavan et al., 2000). Unlike cells isolated from *Itgb1*^+/-^ littermate controls that exhibited increased ICM specification in Matrigel, inside-outside patterning was restored among *Itgb1*^-/-^ cells (Figure 4C). Therefore, ‘inside’ positional signals provided by the ECM require recognition through integrin activity to drive ICM specification.

### Integrin β1 activity is not required for initial specification of ICM but required for EPI-PrE patterning *in vivo*

Earlier observation of integrin β1 mutant mice showed embryonic lethality post-implantation but apparently normal development through preimplantation stages (Fässler and Meyer, 1995). Accordingly, we found TE-ICM patterning and overall morphology to be comparable between wildtype (WT) and *Itgb1*^-/-^ embryos during preimplantion stages (Figure 5A). However, defects were observed upon close examination of late blastocysts. Within the mature ICM, PrE cells form a epithelial monolayer that is apically polarised towards the blastocyst cavity, while EPI cells are sheltered between the PrE and the overlying TE. Although the respective numbers of EPI and PrE cells were not significantly affected by integrin β1 deficiency, PrE cells failed to resolve into a single monolayer in *Itgb1*^-/-^ embryos (Figure 5A and 5B). In addition, distribution of apical PKCζ indicated disrupted polarity in the mutant ICM (Figure 5C and 5D), which, instead of flattening out against the TE, was more rounded in its dimensions (Figure 5E) (Saiz et al., 2013). Therefore, although integrin β1 is not required for initial specification of the ICM *in vivo*, it is required for subsequent spatial organisation among EPI and PrE cells in the blastocyst. These findings reveal that defects that underlie the post-implantation lethality of *Itgb1*^-/-^ embryos in fact arise prior to implantation.

**Figure 5.**
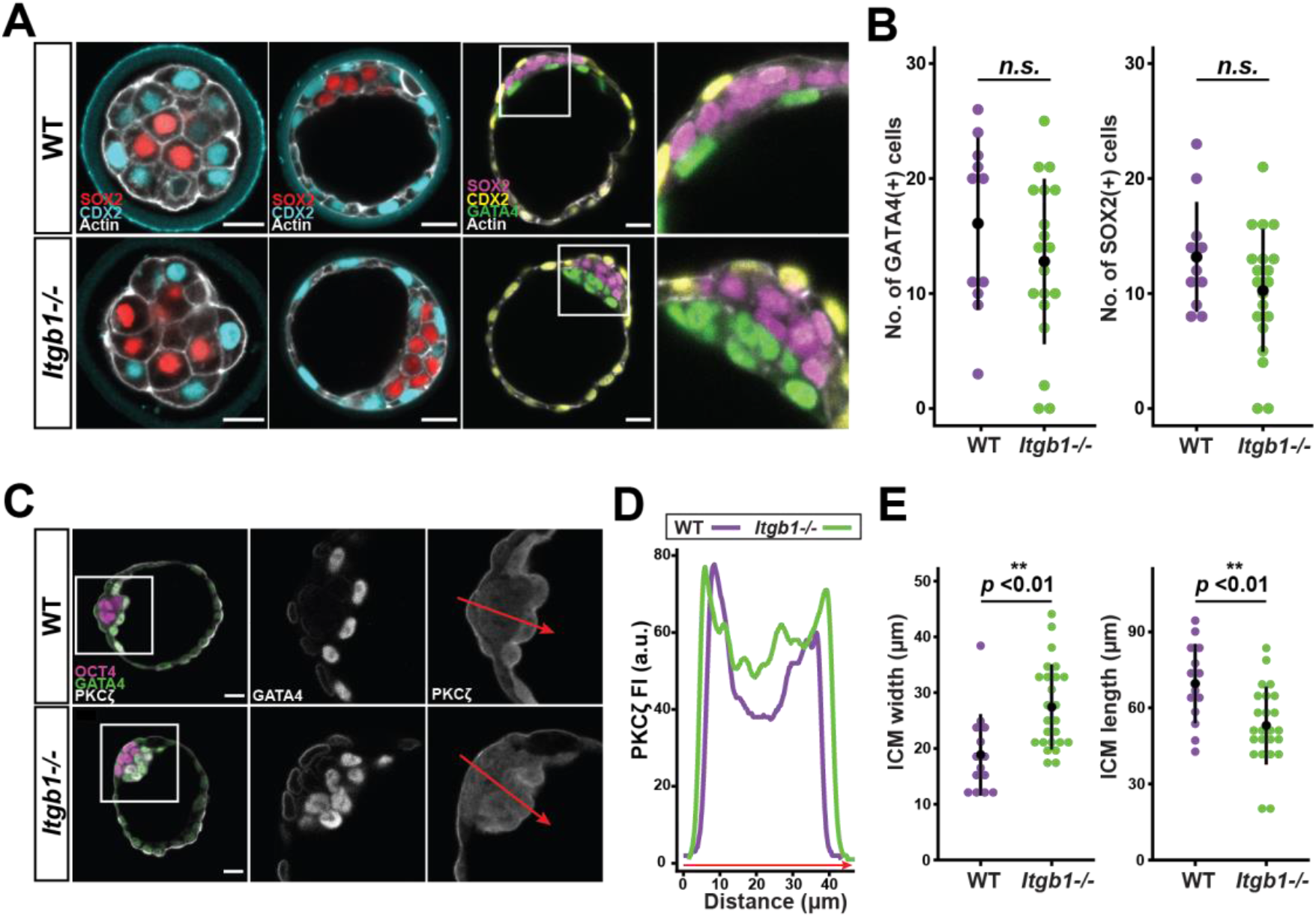
EPI-PrE patterning in the late blastocyst *in vivo* requires integrin β1. (A) Representative images of patterning in preimplantation stage WT and *Itgb1*^-/-^ embryos through morula, early and late blastocyst stages. (B) Cell count of GATA4-expressing PrE cells and SOX2-expressing EPI cells within the ICM of WT and *Itgb1*^-/-^ late blastocysts. Student’s *t*-test, two-sided. Error bars show mean ± s.d. N=31 embryos (11 WT, 20 *Itgb1*^-/-^). (C) Representative image of EPI-PrE arrangement and apical polarity of the ICM of WT and *Itgb1*^-/-^ late blastocysts. OCT4(red) marks EPI, GATA4 (green) marks PrE, and PKCζ(grey) marks apical polarity. (D) Representative intensity profile of PKCζ across the ICM of WT and *Itgb1*^-/-^ late blastocysts along the red line of interest marked in (C). (E) Width and length of the ICM in WT and Itgb1-/- late blastocysts. Student’s *t*-test, two-sided. Error bars show mean ± s.d. N = 39 embryos (14 WT, 25 *Itgb1*^-/-^).

### Exogenous ECM leads to EPI cells dwelling on the surface of the ICM

In contrast to TE-ICM specification, subsequent EPI-PrE specification is not cell position-dependent as respective cells emerge in a salt-and-pepper pattern within the ICM (Chazaud et al., 2006; Plusa et al., 2008). Nevertheless, positional information remains pertinent as EPI and PrE cells must resolve into a distinct spatial pattern as described above. Given the requirement for integrin β1 during this latter process, we tested whether EPI and PrE cells are also receptive to exogenous ECM as positional cues.

Transcription factors NANOG and GATA6 are early markers of EPI and PrE fate, respectively. When ICMs are isolated from blastocyst at E3.5, NANOG- and GATA6-positive nuclei, as well as double-positive nuclei, are intermixed (Figure 6A and 6B). As such, measure of the distance between each nucleus from the centre of the ICM shows no correlation with expression of cell fate markers (Figure 6C).

**Figure 6.**
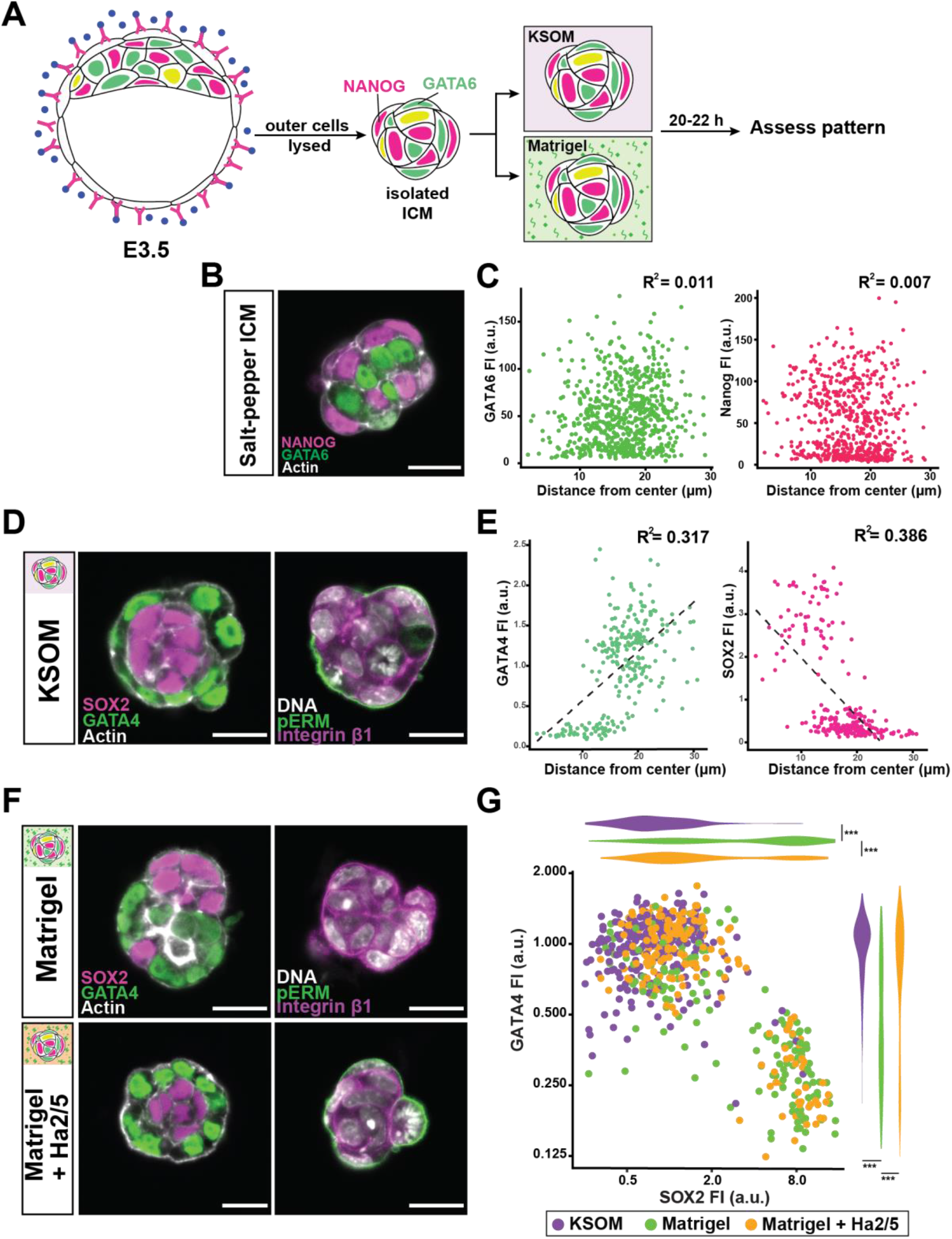
EPI-PrE patterning is sensitive to Matrigel and integrin β1 activity. (A) Schematic representation of experimental conditions with immunosurgery of blastocysts. Blastocysts are subjected to immunosurgery to isolate salt-and-pepper stage ICMs. Isolated ICMs are cultured in either KSOM or Matrigel before assessment of patterning. (B) Representative image of an isolated salt-and-pepper ICM expressing early EPI marker NANOG and early PrE marker GATA6. (C) Scatterplots showing fluorescence intensities of NANOG and GATA6 in relation to cell position within an isolated ICM. Position is measured as distance between each nucleus and the centre of the ICM. Pearson’s correlation. ICMs from N = 28 embryos (n = 658 cells). (D) Representative images of EPI-PrE arrangement and apicobasal polarity of ICMs following culture in KSOM. (E) Scatterplots showing fluorescence intensities of SOX2 and GATA4 in relation to cell position following culture of ICMs in KSOM. Pearson’s correlation. ICMs from N = 35 embryos (n = 765 cells). (F) Representative images of EPI-PrE spatial arrangement and apicobasal polarity of ICMs following culture in Matrigel or Matrigel with integrin β1 function-blocking antibody, Ha2/5 (10μg/ml). Scale bars = 20 μm. (G) Scatterplot and adjacent violin plots show normalised fluorescence intensities of GATA4 and SOX2 measured for each cell cultured in KSOM (purple), Matrigel (green), or Matrigel with Ha2/5 (orange). Mann-Whitney U test. Plot represents data from ICMs from N = 59 embryos (n = 1664 cells). *** *p*< 0.0001

Following immunosurgery and culture, the salt-and-pepper distribution of fates resolved into a pattern where polarised GATA4-positive PrE surrounded the SOX2-positive EPI (Figure 6D). Positional distinction between EPI and PrE was evident based on cell fate marker expression relative to distance from the ICM centre (Figure 6E). In stark contrast, Matrigel markedly disrupted this spatial arrangement (Figure 6F, top left panel). EPI cells were no longer confined to the interior, but frequently found at the surface. Quantitative analysis of fate in peripherally located cells indicated that while the vast majority of outer cells are GATA4-positive in control conditions, a significant portion expresses SOX2 in Matrigel culture (Figure 6G). Furthermore, apical polarity of the ICM surface was replaced by integrin β1 enrichment (Figure 6F, top right panel), as observed from culture of inner cells at the earlier stage.

As with TE-ICM patterning, the effects of Matrigel on EPI-PrE patterning was dependent on integrin β1 activity. Administration of Ha2/5 restored a peripheral polarised PrE layer in the presence of Matrigel (Figure 6F, bottom panel and 6G). These observations demonstrate that ECM-integrin adhesion provides critical positional signals to regulate EPI-PrE patterning within the ICM, consistent with its role in ICM-TE patterning.

### Integrin signalling is mediated by laminin and talin

Given our findings, we next sought to identify extracellular and intracellular components involved in ECM and integrin-mediated position-sensing.

Although laminin itself is a ligand for integrin, integrin β1 is required for the deposition of heterotrimeric laminin, which in turn can bring its cell surface receptors together (Aumailley et al., 2000; Li et al., 2002). Accordingly, intercellular laminin, as judged by strand-like laminin γ1 signals in the ICM, was diminished in *Itgb1*^-/-^ embryos (Figure 7A). Since the requirement for laminin γ1, encoded by *Lamc1*, is shared by both laminin isoforms assembled during the preimplantation stage, our model also predicted integrin signalling to be impaired in *Lamc1*^-/-^ embryos. In fact, PrE cells in *Lamc1*^-/-^ blastocysts failed to resolve into an epithelial monolayer within the ICM (Figure 7B, bottom panels), exactly as observed in *Itgb1*^-/-^ counterparts (Figure 5A). This supports our model in which intercellular laminin provides crucial positional signals that are interpreted by cells via integrin activity to instruct embryonic patterning.

**Figure 7.**
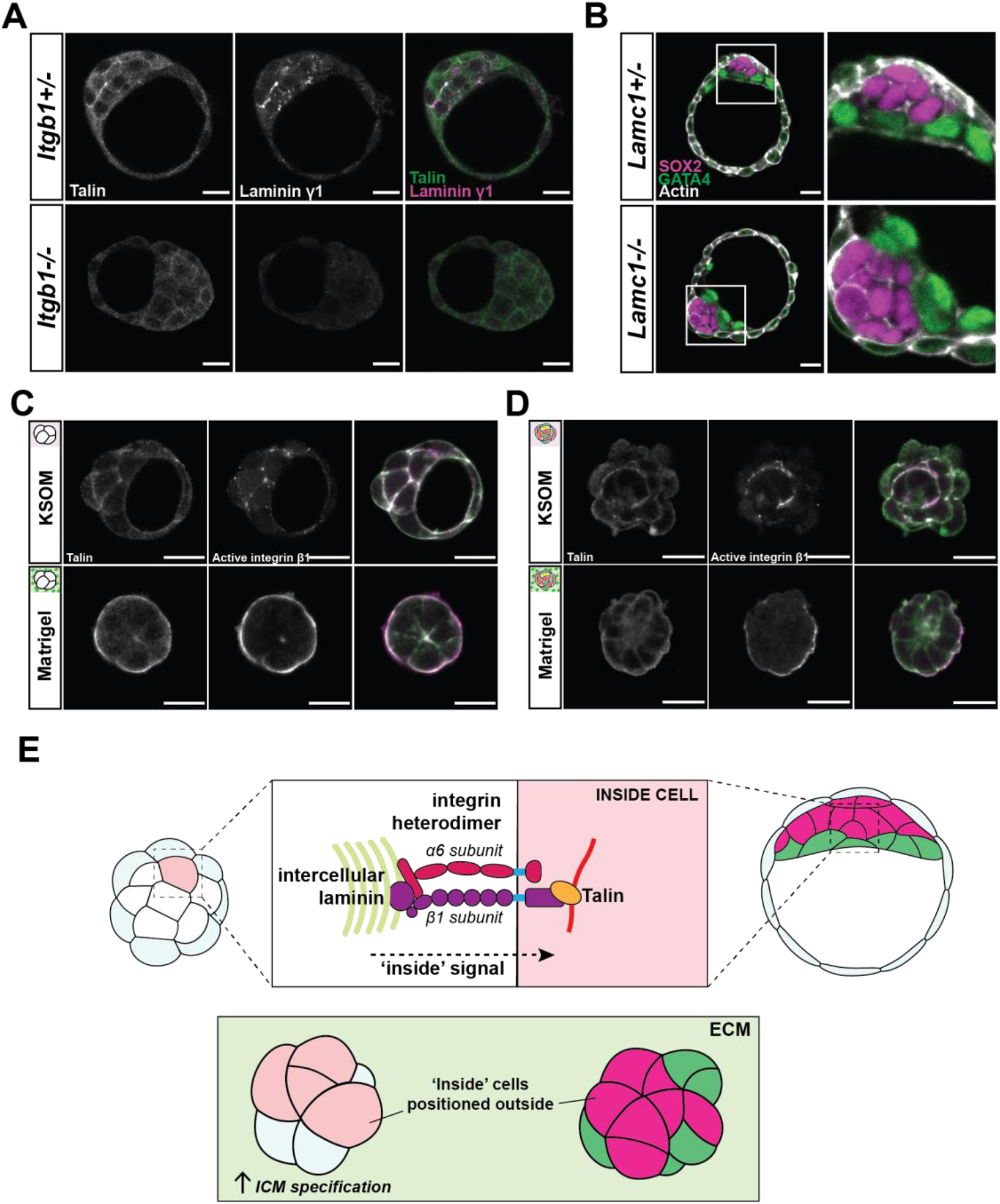
Integrin signalling is mediated by laminin and talin during early embryonic patterning. (A) Representative images of talin and laminin γ1 chain expression in whole blastocysts of *Itgb1*^+/-^ or *Itgb1*^-/-^ genotype. *Itgb1*^+/-^ embryos serve as littermate controls. (B) Representative images of EPI-PrE patterning within the ICM of WT and *Lamc1*^-/-^ late blastocysts. (C and D) Representative images of localisation of talin and the active conformation of integrin β1, following immunosurgery at either E2.5 (C) or E3.5 (D) and culture in KSOM or Matrigel. Scale bars = 20 μm. (E) Schematic outline of laminin-integrin dependent position-sensing by inner cells of the preimplantation mouse embryo. This mechanism for recognition of ‘inside’ position by the cells is shared between morula and blastocyst stage embryos, and modulates cell fate specification as well as spatial arrangement among cells.

Upon ECM ligand binding, the cytoplasmic domain of integrins interacts with a myriad of proteins. Among these, talin plays a key role in linking integrin to the cytoskeleton, and recruits other integrin associated proteins such as vinculin for downstream signalling (Calderwood et al., 1999; Humphries et al., 2007). Indeed, talin is localised to the cell-cell interface in the ICM alongside integrin subunits, and its expression is reduced in *Itgb1*^-/-^ embryos (Figure 7A). Furthermore, where the active conformation of integrin β1 is enriched on the surface by Matrigel culture, talin signal is also increased, both during ICM induction and in surface-positioned EPI cells (Figure 7C and 7D). Together, these strongly suggest that talin functions downstream of integrin to relay positional information from intercellular laminin to early blastomeres located inside the embryo (Figure 7E).

## Discussion

During mouse preimplantation development, ICM-TE specification follows an inside-outside pattern, while EPI and PrE cells initially emerge in an intermixed manner before becoming spatially segregated. Despite this difference, however, we show that cells maintain sensitivity to ECM-integrin signals throughout the preimplantation period to gain positional information. Given that altered patterning induced by Matrigel required integrin α6β1 activity, it follows that laminin, rather than other factors associated with reconstituted ECM, are pertinent for patterning the early embryo.

In developing embryos or stem cell systems where cells are yet to differentiate, a myriad of signals must be processed leading up to lineage commitment. During the first lineage segregation, Matrigel is sufficient to drive ICM specification in an integrin-dependent manner, irrespective of cell position. Yet, integrin activity is not required for inside-outside patterning *per se*. Given the significance of setting aside cells that will eventually form the embryo proper, other factors, such as the non-integrin laminin receptor dystroglycan, may well be active in the embryonic interior as redundant ‘inside’ signals (Hynes, 1987; Mui et al., 2016; Williamson et al., 1997). In addition, the ECM provided through Matrigel in our setup may be more concentrated than levels found *in vivo*, thereby overriding competing positional signals to drive ICM specification.

Our work complements earlier studies in embryonic stem cells that revealed ECM-integrin signals as critical regulators of the undifferentiated state and cell arrangement (Aumailley et al., 2000; Cattavarayane et al., 2015; Li et al., 2002). Given the ubiquity and tissue/stage-dependent complexity of the ECM and its receptors, their role in cell fate specification and pattern formation extends beyond early mouse development (Huang and Ingber, 2005; Humphrey et al., 2014; Walma and K. M. Yamada, 2020; Watt and Huck, 2013). Elucidation of their mechanistic contribution to patterning across diverse contexts will be instrumental to how we approach various disease states and design regenerative therapies in the future.

## Materials & Methods

### Animal work

All animal work was performed in the Laboratory Animals Resources (LAR) facility at the European Molecular Biology Laboratory (EMBL) with permission from the Institutional Animal Care and Use Committee (IACUC) overseeing the operations (IACUC #TH110011). The LAR facility operates according to guidelines and recommendations set by the Federation for Laboratory Animal Science Associations. Mice were maintained in pathogen-free conditions under 12-hour light-dark cycles.

#### Mouse lines

Wildtype mice were of a F1 hybrid strain from C57BL/6 and C3H (B6C3F1) animals. The following transgenic lines were used in this study: Sox2-GFP (Arnold et al., 2011), mTmG (Muzumdar et al., 2007), *Itgb1^tm1Efu (floxed)^* (Raghavan et al., 2000), *Lamc1^tmStrl(floxed)^* (Chen and Strickland, 2003), Zp3-Cre (de Vries et al., 2000). Standard tail genotyping procedures were used to genotype transgenic mice.

To obtain Sox2-GFP x mTmG embryos, Sox2-GFP animals were crossed with mTmG animals. To obtain *Itgb1*^+/-^ mice, *Itgb1^tm1Efu (floxed)^ Zp3-Cre^tg^* females were crossed with B6C3F1 males. To obtain zygotic *Itgb1*^-/-^ embryos, *Itgb1*^+/-^ females were crossed with *Itgb1*^+/-^ males. To obtain *Lamc1*^+/-^ mice, *Lamc1^tmStrl (floxed)^ Zp3-Cre^tg^* females were crossed with B6C3F1 males. To obtain zygotic *Lamc1*^-/-^ embryos, *Lamc1*^+/-^ females were crossed with *Lamc1*^+/-^ males.

#### Superovulation and dissection of reproductive organs

Superovulation was induced in females prior to mating to increase the number of preimplantation embryos obtained per mouse. Intraperitoneal injection of 5IU of PMSG (Intervet, Intergonan) and hCG (Intervet, Ovogest 1500) were carried out, with a 48-50 hour interval between the two injections. Each female mouse was put in a cage with a male immediately following hCG injection for mating.

Timing of sacrifice post-hCG injection depends on the developmental stage relevant for the experiment. Given 11AM hormone injections for superovulation, 16-32 cell stage embryos were recovered in the afternoon of E2.5. Early blastocysts were obtained on E3.5.

### Embryo work

Preimplantation embryos were obtained by flushing the oviduct with a 1ml syringe filled with H-KSOM from the infundibulum. All live embryos were handled under a stereomicroscope (Zeiss, Discovery.v8) equipped with a heating plate (Tokai hit, MATS-UST2). All live embryos were cultured in 10μl microdroplets of KSOM (potassium Simplex Optimized Medium; (Lawitts and Biggers, 1991)) with a mineral oil (Sigma, M8410) overlay inside an incubator (Thermo Fisher Scientific, Heracell 240i) with a 37°C humidified atmosphere of 5% CO2 and 95% air. Micromanipulations outside the incubator were carried out in KSOM containing HEPES (H-KSOM; LifeGlobal, LGGH-050).

#### Immunosurgery

Zona pellucida were removed from embryos with 3-4 min pronase (0.5% w/v Proteinase K, Sigma P8811, in H-KSOM supplemented with 0.5% PVP-40) treatment at 37°C Subsequently, embryos were incubated in serum containing anti-mouse antibody (Cedarlane, CL2301, Lot no. 049M4847V) diluted 1:3 with KSOM for 30 min at 37°C. Following three brief washes in H-KSOM, embryos were incubated in guinea pig complement (Sigma, 1639, Lot no. SLBX9353) diluted 1:3 with KSOM for 30 min at 37°C. Lysed outer cells were removed by mouth-pipetting with a narrow glass capillary to isolate the inner cells.

#### Embedding cells in Matrigel

Matrigel mix consists of Matrigel (Corning 356230, lot. 7345012) diluted in DPBS to desired concentration (4.5 mg/ml). Matrigel mix was prepared fresh for each experiment, mixed thoroughly through pipetting, and kept on ice during immunosurgery. Upon completion of immunosurgery, isolated inner cells were promptly resuspended in the mixes, and 15μL droplets were made on 35mm petri dishes (Falcon, 351008). To ensure that cell clusters from different embryos do not stick together, a closed glass capillary was used to space them apart. These petri dishes were inverted to prevent cells sticking to the bottom of the dish, and incubated at 37°C for 30 minutes for the mix to form a gel. After gel formation, 4ml of prewarmed KSOM was gently pipetted into each dish to cover the gel.

To inhibit integrin heterodimer activity in Matrigel-embedded cells, blocking antibodies Ha2/5 and GoH3 that target integrin β1 and α6, respectively, were added to the overlying KSOM medium at a concentration of 10 μg/mL.

#### Immunostaining

Embryos were fixed in 4% PFA (Sigma, P6148) at room temperature for 15 min, washed 3 times (5 min each) in wash buffer (DPBS-T containing 2% BSA), permeabilised at room temperature for 30 min in permeabilisation buffer (0.5% Triton-X in DPBS; Sigma T8787), washed (3 x 5 min), followed by incubation in blocking buffer (PBS-T containing 5% BSA) either overnight at 4°C or for 2 h at room temperature. Blocked samples were incubated with primary antibodies (Table1) overnight at 4°C, washed (3 x 5 min), and incubated in fluorophore-conjugated secondary antibodies and dyes (**Error! Reference source not found.**) for 2 hours at room temperature. Stained samples were washed (3 x 5 min), incubated in DAPI solution (Life Technologies, D3571; diluted 1:1000 in DPBS) for 10 min at room temperature. These samples were then transferred into droplets of DPBS overlaid with mineral oil on a 35mm glass bottom dish (MatTek, P356-1.5-20-C) for imaging.

### Molecular work

#### Single embryo genotyping

Individual embryos were mouth pipetted into 200μL PCR tubes containing 10μL of lysis buffer consisting of 200μg/ml Proteinase K in *Taq* polymerase buffer (Thermo Scientific, B38). Lysis reaction took place for 1 hour at 55°C, followed by 10 minutes at 96°C. Resulting genomic DNA was mixed with relevant primers (Table 3) for determination of genotype via PCR

**Table 1.** Primary antibodies.

| **Epitope** | **Host** | **Catalogue #** | **Company** | **Dilution** |
| --- | --- | --- | --- | --- |
| aPKC (PKCζ) | Rabbit | sc-216 | Santa Cruz Biotechnology | 1:200 |
| CDX2 | Mouse | MU392A-UC | Biogenex | 1:200 |
| E-cadherin | Rat | U3254 | Sigma Aldrich | 1:100 |
| GATA4 | Goat | AF2606 | R&D Systems | 1:200 |
| GATA6 | Goat | AF1700 | R&D Systems | 1:200 |
| Integrin α6<br>(GoH3) | Rat | 555734 | BD Pharmingen | 1:100 |
| Integrin β1 | Rat | MAB1997 | Millipore | 1:100 |
| Integrin β1<br>(Ha2/5) | Rat | 555002 | BD Pharmingen | 1:100 |
| Laminin (non-chain specific) | Rabbit | NB300-14422 | Novus Biologicals | 1:100 |
| Laminin α5 | Rat | n/a | Gift from Lydia Sorokin | N/A |
| Laminin β1 | Rat | n/a | Gift from Lydia Sorokin | N/A |
| Laminin γ1 | Rat | n/a | Gift from Lydia Sorokin | N/A |
| NANOG | Rabbit | RCAB002P-F | ReproCELL, Inc | 1:200 |
| YAP1 | Mouse | H00010413-M01 | Abnova | 1:100 |
| Phospho-ERM | Rabbit | 3726 | Cell Signaling Technology | 1:200 |
| Sox-2 (D9B8N) | Rabbit | 23064 | Cell Signaling Technology | 1:200 |

**Table 2.** Secondary antibodies and dyes.

| **Fluorophore** | **Target** | **Host** | **Catalogue #** | **Company** | **Dilution** |
| --- | --- | --- | --- | --- | --- |
| Alexa Fluor 488 | Goat IgG | Donkey | A11055 | Life Technologies | 1:200 |
| Alexa Fluor 488 Plus | Rabbit IgG | Donkey | A32790 | ThermoFisher | 1:200 |
| Alexa Fluor 546 | Rabbit IgG | Donkey | A10040 | ThermoFisher |  |
| Cy5 | Mouse IgG | Donkey | 715-175-150 | Jackson<br>ImmunoResearch | 1:200 |
| Cy5 | Rat IgG | Donkey | 712-175-153 | Jackson<br>ImmunoResearch | 1:200 |
| DAPI | (DNA) | - | D3571 | Life Technologies | 1:1000 |
| Rhodamine phalloidin | (Actin) | - | R415 | Invitrogen | 1:200 |

**Table 3.**
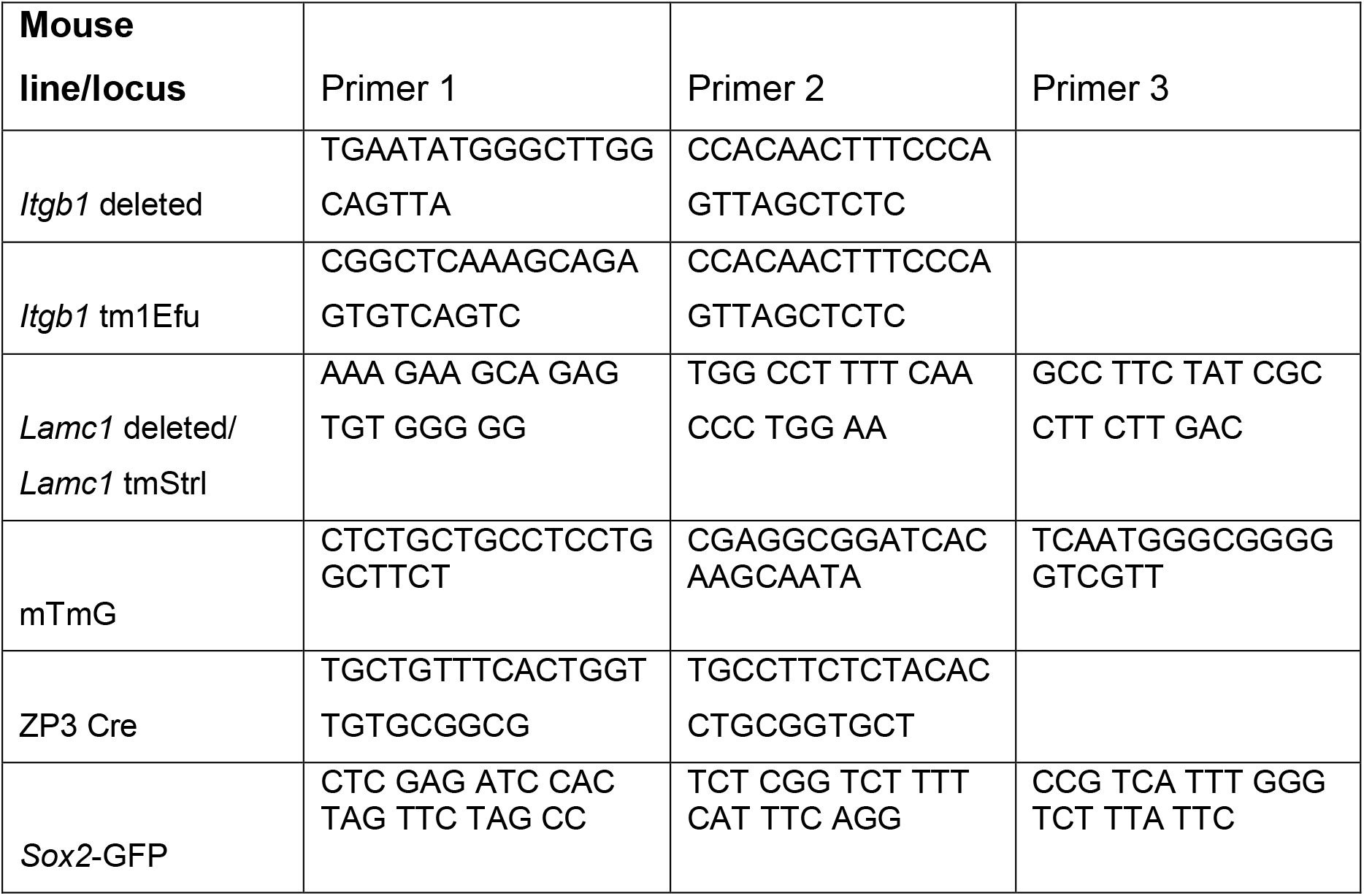
Sequence of genotyping primers.

#### Microscopy and image analyses

Fixed and stained embryos were imaged on the Zeiss LSM780 and LSM880 confocal microscopes. For both systems, a 40X water-immersion C-Apochromat 1.2 NA objective lens was used. Imaging was carried out with the Zen (Zeiss) software interface. Resulting raw images were processed using ImageJ. Further quantification of fluorescence intensities and nuclei/cell counting was performed on either ImageJ or Imaris 9.2.1 (Oxford Instruments) as described below.

#### Light-sheet live-imaging

*Sox2*-GFP x mT embryos were live-imaged using InVi SPIM (Strnad et al., 2016). Embryos were mounted on a V-shaped sample holder covered with transparent FEP foil. Each embryo was placed in a small microwell made by gently pressing a closed glass pipette into the foil. Embryos were immersed in 100 μL KSOM medium and covered in 150 μL mineral oil to prevent evaporation. The samples were imaged inside an environmentally controlled incubation chamber with 5% CO_2_ and 5% O_2_ at 37 °C.

InVi SPIM was equipped with a Nikon 25x/1.1NA water immersion detective objective and a Nikon 10x/0.3 NA water immersion illumination objective. Illumination plane and focal plane were aligned before each imaging session. Images were taken every 15 minutes from the 8-cell stage to the early blastocyst stage by a CMOS camera (Hamamatsu, ORCA Flash4.0 V2) with line-scan mode.

#### Quantification of *Sox2*-GFP fluorescence intensity during ICM specification

GFP fluorescence intensity was measured at each time point by drawing a region of interest (ROI) within the cross-section of each ICM cell and obtaining the mean grey value on ImageJ. Each ICM cell was identified in the blastocyst based on GFP fluorescence and retrospectively tracked to identify the point of internalisation. A cell was considered to be ‘internalised’ when its entire surface was surrounded by adjacent cells, with no area exposed to the external environment. Tracking extended to parent cells prior to *Sox2* upregulation. For each embryo, 8-11 ICM cells were tracked in this way.

Background fluorescence was quantified by calculating the average intensity in blastomeres during the 8-cell, morula, and blastocyst stages. This value was subtracted from measured *Sox2*-GFP intensity of ICM cells.

#### Measure of cell circularity

Circularity measurement were obtained by tracing the outline of individual cells on ImageJ. From each cluster, up to 4-8 TE- and ICM-specified cells were traced across their mid-section. Cells undergoing division were not included, as cells round up during mitosis.

#### Measure of ICM width and length in blastocysts

A confocal image slice was taken through the mid-section of the ICM. Width was defined as the distance between the centre of the PrE layer and the ICM-TE boundary. Length was defined as the distance between two PrE cells at the opposite ends. A straight line was drawn for width and length, and the distance was measured in ImageJ.

#### Quantification of fluorescence intensity of lineage markers

For measure of lineage specification, Imaris was used. Imaris Surpass allowed 3D visualisation of the image data. The ‘Add Spots’ function was used to detect each nucleus on the DAPI channel. Estimated spot (nucleus) diameter was set to 6 μm, and manual corrections were made for each image as necessary to detect all nuclei. Mean fluorescence intensity of SOX2, CDX2 or GATA4 were measured for each detected nucleus. Spot detection of each nucleus was also used as a cell counter.

#### Quantification of fluorescence intensity of apicobasal markers

Fluorescence signal intensity of cortical pERM and integrin β1 was used as a measure of apical and basal polarity, respectively. Images of samples stained for these proteins were analysed on ImageJ. To reduce noise, the Gaussian filter was applied to smooth the image. For each z-stack, a mid-slice was selected, and a line was traced along the perimeter of the smoothed image. A plot profile along the line was obtained for the pERM and integrin β1 channels. Individual data points were exported from ImageJ for statistical analysis.

#### Statistical analysis

Statistical analyses and graph generation was performed using the ggplot2 package in R and Microsoft Excel. Comparison of the distribution of fate marker intensities was performed by the Mann-Whitney U test. Differences in cell count, surface enrichment of apicobasal polarity markers, circularity, and ICM dimensions were assessed using the student’s *t*-test (two-sided). Statistical dependence between *Sox2*-GFP intensity and duration a cell resides in the embryonic interior, and relationship between EPI/PrE fate marker expression and cell position were assessed by Pearson’s correlation.

## Acknowledgements

We are grateful to members of the Hiiragi Group and Aissam Ikmi for critical comments and suggestions. Karen-Sue Carlson kindly provided *Lamc1 ^fl/fl^* mice. We also thank Ramona Bloehs, Stefanie Friese, Lidia Perez, and the Laboratory Animal Resources at EMBL for technical support with mouse work. E.J.Y.K. is supported by the EMBL International PhD Programme, and the Hiiragi Group is supported by EMBL and the European Research Council (ERC Advanced Grant “Self-organising Embryo”, grant agreement 742732).

## Author contributions

The project was conceived and designed by E.J.Y.K and T.H. L.S. provided expertise and reagents relating to the ECM, particularly laminin, which further guided the design of the project. E.J.Y.K. carried out all experiments, including mouse experiments, imaging and data analysis under the supervision of T.H. E.J.Y.K. wrote the manuscript, which was reviewed and edited by L.S. and T.H. for the final version. T.H. acquired the funding.

---

**Figure S1.**
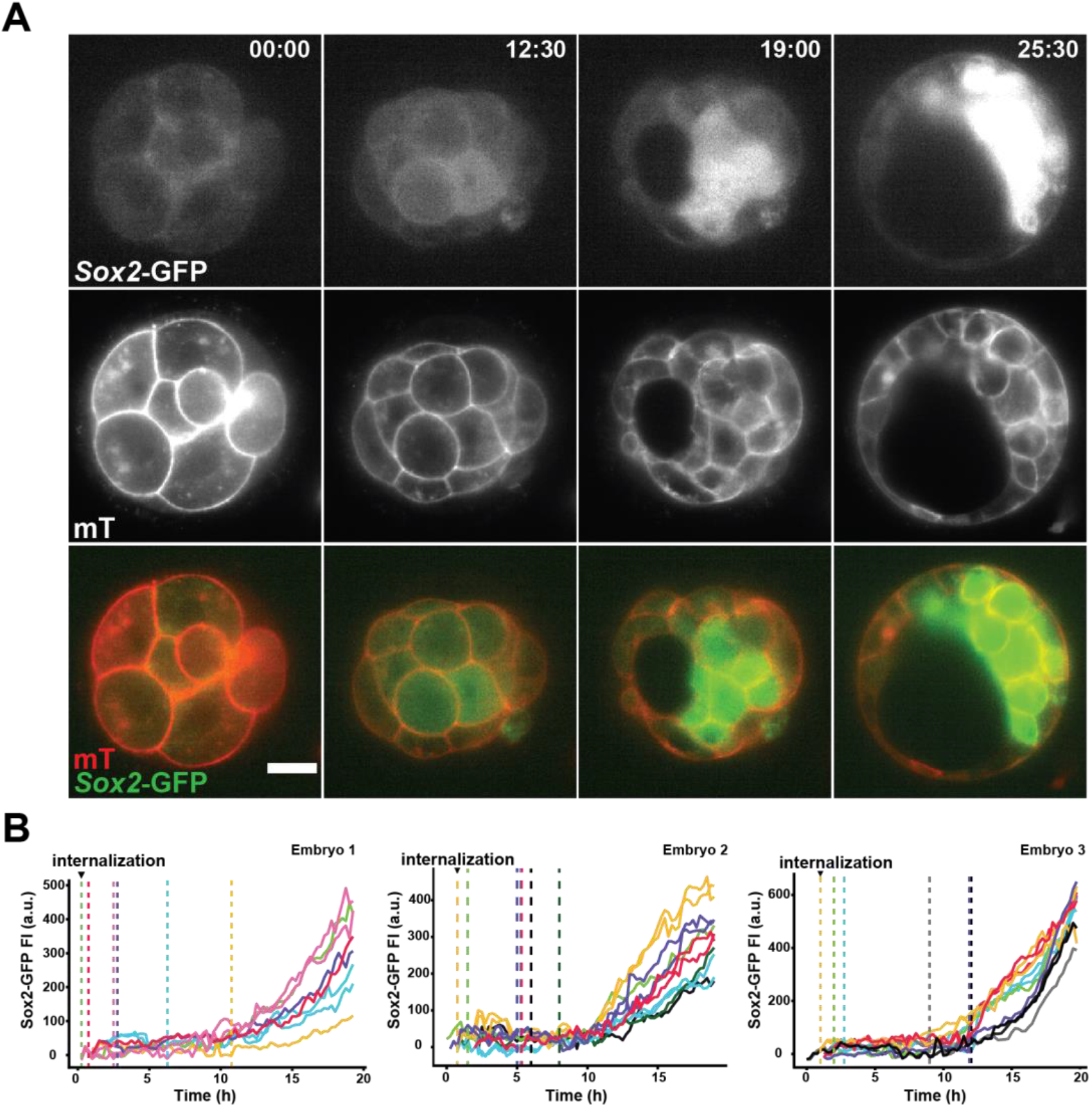
ICM fate specification is induced by inner positioning of blastomeres. Related to Figure 1. (A) Time-lapse images of ICM specification during preimplantation development of a *Sox2*-GFP x mT embryo. Time, post 8-to-9 cell division (h:min). Scale bar = 20 μm. (B) Changes in *Sox2*-GFP intensity during ICM specification in different embryos. For each blastocyst, 8-11 SOX2-positive cells were retrospectively tracked and GFP intensities were measured. Cells become internalised at different timepoints (vertical dashed lines). Each line represents a single cell that becomes ICM-specified, and sister cells are marked in the same colour.

**Figure S2.**
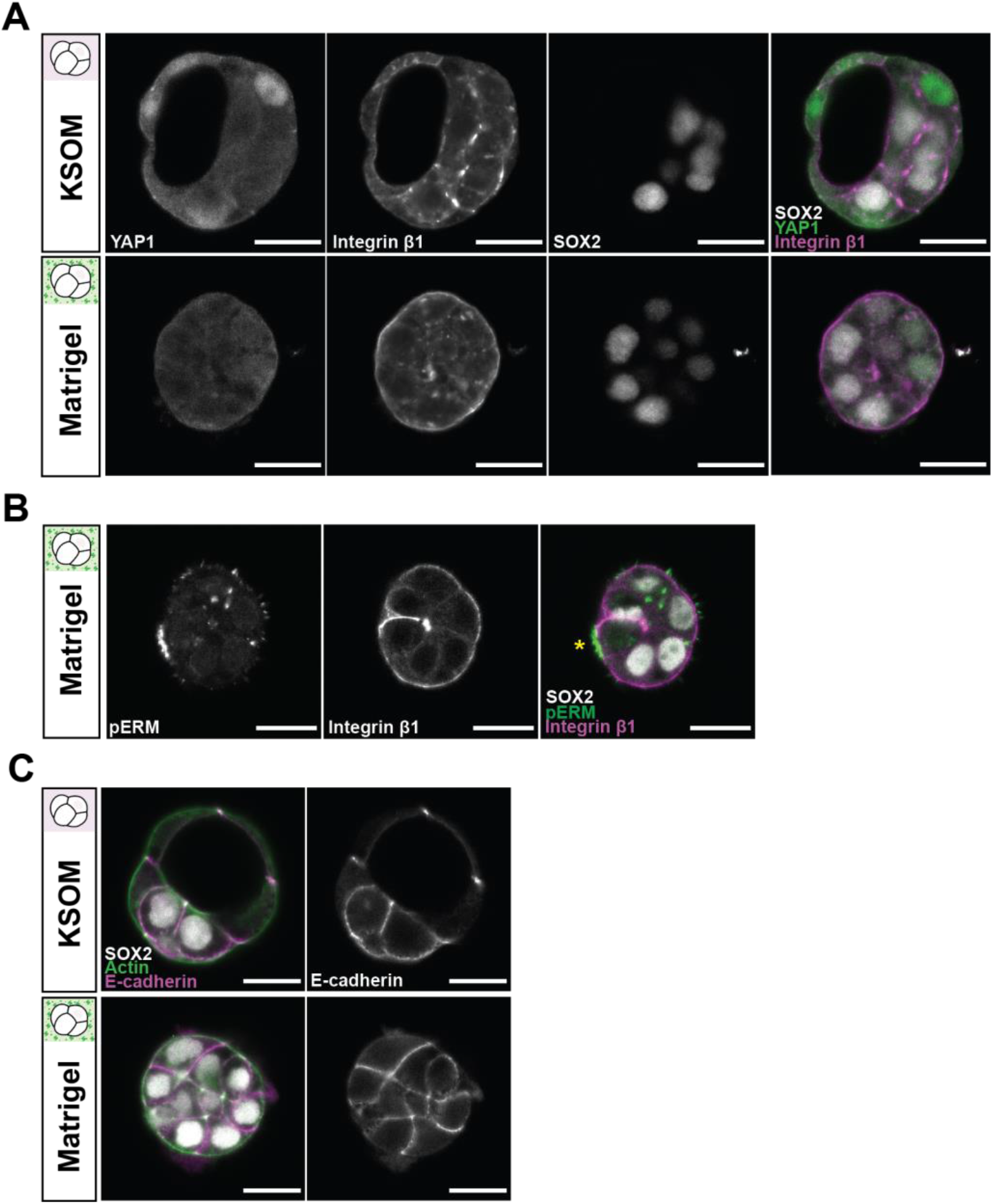
Exogenous ECM drives Hippo signalling and suppresses apical polarity. Related to Figure 3. (A) Representative images of TE-ICM fate specification in inner cells following immunosurgery and culture in KSOM or Matrigel. SOX2 marks ICM fate while nuclear YAP1 is characteristic of TE cells. (B) Partial enrichment of pERM on the surface of isolated cells cultured in Matrigel. The cell with the patch of pERM signal (*) is SOX2-negative. (C) Representative images of E-cadherin localisation in isolated cells following culture in KSOM or Matrigel. Scale bar = 20 μm.

**Figure S3.**
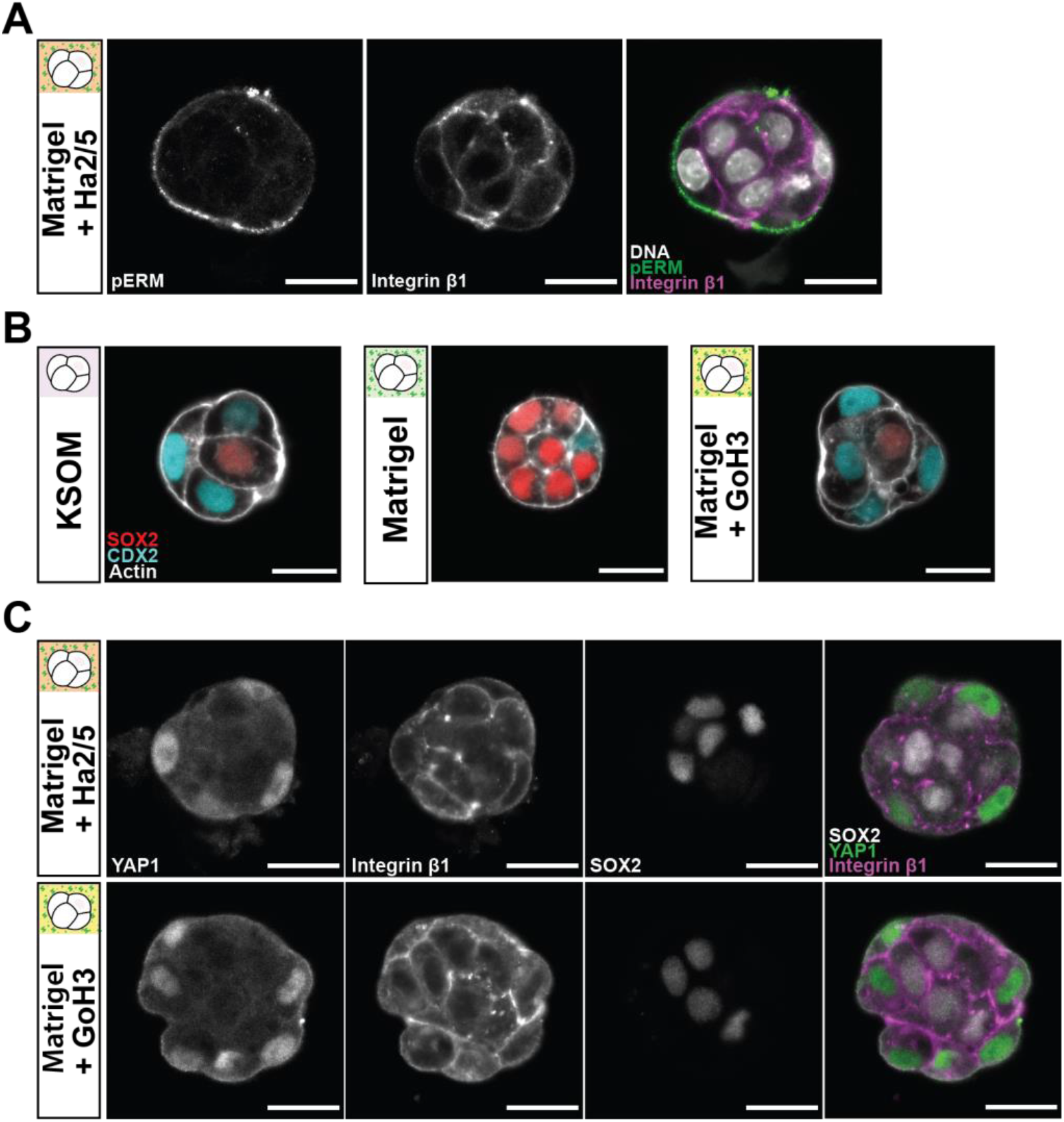
Integrin α6β1 inhibition restores inside-outside patterning to Matrigel-cultured cells. Related to Figure 4. (A) Representative images of apicobasal polarity in cells cultured in Matrigel with integrin β1 function-blocking antibody Ha2/5 (10 μg/ml). Phosphorylated ERM (pERM) proteins mark apical domain. (B) Representative images of TE-ICM fate specification following culture in KSOM, Matrigel, or Matrigel with integrin α6 function-blocking antibody GoH3 (10 μg/ml). (C) Representative images of inside-outside patterning following culture in Matrigel with either Ha2/5 or GoH3. In addition to SOX2 expression, differential localisation of YAP1 distinguishes TE and ICM fate, as YAP1 is nuclear localised in TE cells.

**Movie S1 and S2.** Time-lapse videos of ICM specification during preimplantation development of a representative *Sox2*-GFP x mT embryos. Movie S1 corresponds to Figure 1B. Movie S2 corresponds to Figure S1A. Time, post 8-to-9 cell division (h:min).

